# Marker subset selection and decision support range identification for acute myeloid leukemia classification model development with multiparameter flow cytometry

**DOI:** 10.1101/621797

**Authors:** Jang-Sik Choi, Nguyen Thanh Nguyen, Hyung-Gi Byun, Jaewoo Song, Tae-Hyun Yoon

## Abstract

In this study, we developed acute myeloid leukemia (AML) classification model through Wilks’ lambda-based important marker-identification method and stepwise–forward selection approach, and spotted important decision-support range of flow-cytometry parameter using insights provided by machine-learning algorithm. AML flow-cytometry data released from FlowCAP-II challenge in 2011 was used. In FlowCAP-II challenge, several sample classification algorithms were able to effectively classify AML and non-AML. Most algorithms extracted features from high-dimensional flow-cytometry readout comprised of multiple fluorescent parameters for a large number of antibodies. Multiple parameters with forward scatter and side scatter increase computational complexity in the feature-extraction procedure as well as in the model development. Parameter-subset selection can decrease model complexity, improve model performance, and contribute to a panel design specific for target disease. With this motivation, we estimated importance of each parameter via Wilks’ lambda and then identified the best subset of parameters using stepwise–forward selection. In the importance-estimation process, histogram matrix of each parameter was used. As a result, parameters, which are associated with blasts gating and identification of immature myeloid cells, were identified as important descriptors in AML classification, and combination of these markers is more effective than an individual marker. A random-forest, supervised-classification machine-learning algorithm was used for the model development. We highlighted decision-support range of the fluorescent signal for the identified important parameters, which significantly contribute to AML classification, through a mean decrease in Gini supported in random forest. These specific ranges could help with establishing diagnosis criteria and elaborate the AML classification model. Because methodology proposed in this study can not only estimate the importance of each parameter but also identify the best subset and the specific ranges, we expect that it would contribute to *in silico* modeling using flow- and mass-cytometry readout as well as panel design for sample classification.

**Author summary:** Flow cytometry is a widely used technique to analyze multiple physical characteristics of an individual cell and diagnose and monitor human disease as well as response to therapy. Recent developments in hardware (multiple lasers and fluorescence detectors), fluorochromes, and antibodies have facilitated the comprehensive and in-depth analysis of high numbers of cells on a single cell level and led to the creation of various computational analysis methods for cell type identification, rare cell identification, and sample classification. Flow cytometry typically uses panels with a large number of antibodies, leading to high-dimensional multiparameter flow cytometry readout. It increases computational complexity and makes interpretation difficult. In this study, we identified the best subset of the parameters for AML classification model development. The subset would contribute to panel design specific for the target disease and lead to easy interpretation of the results. In addition, we spotted important decision-support range of flow-cytometry parameter via insights provided by machine-learning algorithm. We expect that profiling information of fluorescence expression over the identified decision-support range would complement existing diagnosis criteria.

## Introduction

Acute myeloid leukemia (AML) is an aggressive, rapidly fatal blood cancer, characterized by the rapid growth of abnormal immature myeloid cells in both the bone marrow and peripheral blood [1]. Because AML is a heterogeneous disease with several subtypes, the prognosis and treatment depend on the subtype [2,3].

Flow cytometry has become an essential method for diagnosis and identification of AML because of its excellent auxiliary diagnostic performance [4–8]. It is a bioanalytical technique to measure and analyze multiple physical and chemical characteristics of tens of thousands of cells via a laser light interacting with monoclonal antibodies specific for a cell surface antigen [9]. The use of multiple lasers and a wide range of antibodies allows clinicians to make specific diagnoses and classifications of various hematological diseases [10]. A typical sample such as blood, bone-marrow tissue, body fluids, and lymph nodes is usually stained with dozens of monoclonal antibodies. Depending on the capabilities of the flow cytometry instrument, 3–10 or more single-tube combinations of the antibodies per sample are used to characterize different cellular populations.

In flow cytometry, traditional data-analysis methods rely on subjective manual gating, which draws regions of interest (gates) on a plot, identifies subsets, and estimates statistics of the subsets (e.g., relative proportion, median fluorescence intensity (MFI), etc.). Because flow cytometry typically uses panels with a large number of antibodies, leading to high-dimensional data, manual gating has been considered as a time-consuming and labor-intensive process [11]. In addition, the manual-gating approach relies on the researcher’s prior knowledge, thus introducing a bias toward “expected" results [12]. Therefore, there has been high demand for the development and application of *in silico* computational methods to flow-cytometry data to overcome these problems in the manual gating-based data-analysis method.

In 2011, FlowCAP (Flow Cytometry: Critical Assessment of Population Identification Methods) challenges were established to compare the performance of computational methods on two tasks: (i) cell-population identification and (ii) sample classification (http://flowcap.flowsite.org/) [13]. In the sample-classification challenges, several classification algorithms were able to effectively classify AML and non-AML samples. Majority of the algorithms extracted a feature, which represents the high-dimensional flow-cytometry data of each sample, through clustering methods and 2D histogram computation for the classification-model development. Because of various markers used in AML samples, the feature was extracted from flow-cytometry data comprised of multiple fluorescent parameters in a sample. The multiple fluorescent parameters increase computational complexity in the whole process of model development. Parameter-subset selection not only can decrease the complexity but also can lead to better performance of the model. In addition, it could contribute to a panel design specific for a target disease. It motivated our study to estimate the importance of each marker and identify the best subset of these markers for the optimization of AML classification model.

The AML classification problem of identifying to which set of categories (AML or non-AML) a new sample belongs can be modelled using machine-learning algorithms. Machine-learning algorithms have been used in a wide variety of fields and also applied at various stages such as feature extraction, feature-importance estimation, and data-dimension reduction when developing a classification model. In particular, feature-importance estimation allows identification of essential features, which plays an important role in sample classification. It can provide new insights on a classification model using multiparameter flow-cytometry data when features, which represent event expression change over the range of each fluorescent channel signal for a specific antibody, are used as input for a machine-learning algorithm-based classification model. With this benefit of machine-learning algorithms, we identified a specific decision-support range of the fluorescent signal for the identified important parameters, using feature importance estimated from a machine-learning model.

In this study, we computed the importance of each parameter using Wilks’ lambda (Λ) and then used stepwise–forward selection method to select the best subset of parameters. In addition, we identified decision-support range of the fluorescent signal for the identified important parameters, which significantly contribute to AML classification, through a mean decrease in Gini (MDG) supported in a random-forest model. We expect that the methodology presented in this study could contribute to *in silico* sample-classification modelling using flow- or mass-cytometry readout because it can provide the importance of each parameter, the best subset of parameters, and the specific ranges in terms of sample discrimination.

## Materials and methods

### Data

AML data released from FlowCAP-II challenge in 2011 was used in this study. The dataset contains 2872 flow-cytometry standard (FCS) files obtained from peripheral-blood or bone-marrow samples of 359 patients (AML patients = 43, healthy patients = 316). Because of the large number of parameters used, eight tubes with different combinations of fluorescent markers (reagents) were used. Therefore, there are eight FCS files per sample in the data. The parameters (reagents and channels) used in each tube are listed in Table 1. CD45-ECD, forward scatter (FSC), and side scatter (SSC) are available in all tubes. In this study, the data were logarithmically transformed for SSC and all fluorescent parameters, while FCS data remained linear. The values of all parameters range from 0 to 1024. The same data labels used in FlowCAP-II challenge were applied for model training and testing in this study. In the challenge, data was split into a training set (179 patients) and testing set (180 patients). The training set includes 156 healthy patients and 23 AML patients, while the testing set has 160 and 20 normal and AML patients, respectively. The FlowCAP-II AML data is publicly available at the flow repository: http://flowrepository.org/id/FR-FCM-ZZYA.

**Table 1.** Eight tubes with different fluorescent reagent combination.

| No. | Tube |  |  |  |  |  |  |  |
| --- | --- | --- | --- | --- | --- | --- | --- | --- |
|  | 1 | 2 | 3 | 4 | 5 | 6 | 7 | 8 |
| 1 | FSC | FSC | FSC | FSC | FSC | FSC | FSC | FSC |
| 2 | SSC | SSC | SSC | SSC | SSC | SSC | SSC | SSC |
| 3 | IgG1-FITC | Kappa-FITC | CD7-FITC | CD15-FITC | CD14-FITC | HLA DR-FITC | CD5-FITC | FL1 |
| 4 | IgG1-PE | Lambda-PE | CD4-PE | CD13-PE | CD11c-PE | CD117-PE | CD19-PE | FL2 |
| 5 | CD45-ECD | CD45-ECD | CD45-ECD | CD45-ECD | CD45-ECD | CD45-ECD | CD45-ECD | 7AAD |
| 6 | IgG1-PC5 | CD19-PC5 | CD8-PC5 | CD16-PC5 | CD64-PC5 | CD34-PC5 | CD3-PC5 | FL4 |
| 7 | IgG1-PC7 | CD20-PC7 | CD2-PC7 | CD56-PC7 | CD33-PC7 | CD38-PC7 | CD10-PC7 | FL5 |

### Methods

#### Feature extraction

Each FCS file contains thousands of events (rows) and 7 features (columns) corresponding to 7 fluorescent channels of each tube. The events are measurements for a series of cells. In FlowCAP-II challenge, a feature vector, which represents the high-dimensional data of FCS file, was extracted using manual gating, auto gating, and 2D histograms of all possible pairs of parameters. The feature vector was comprised of statistical values extracted from clusters identified by the gating methods or by the proportion of cells that fall into a specific bin on a 2-dimensional scatter plot. Manual gating is effective but subjective and extremely laborious. Auto gating using clustering algorithms requires high computational cost and expert knowledge to identify cell population based on the results. In addition, clustering results are dependent on coordinates of an initial random cluster, distance measure, and number of clusters. The 2D histogram-based approach yields a number of combinations from *n* parameters if a large number of markers are used.

In this study, 1D histogram, which displays a single measurement parameter (fluorescence intensity) on the x-axis and the number of events (cell count) on the y-axis, was used as a feature vector of each parameter because it allows exploration on expression change over the range of each fluorescent channel signal for a specific antibody. The histogram used in this study has equal-sized 32 bins in the range 0 to 1024. Two or more histograms for parameters in the FCS sample file were concatenated into a single histogram vector. Thereby, the high-dimensional data of an FCS file was represented by a single histogram vector.

#### Important marker selection

Measuring the importance of each parameter allows identification of the best combination of parameters. In addition, feature selection based on importance is capable of improving learning performance, lowering computational complexity, building a better generalizable model, and decreasing required storage [14]. To measure the importance of each parameter, we used Wilks’ lambda (Λ), which is a measure of how well each independent parameter contributes to the group separation [15,16]. A smaller Λ value indicates greater discriminatory ability of the related parameter.

Histogram matrix for training data (179 patients) was used to measure Λ values of a total of 56 parameters (8 tubes and 7 channels). For example, Λ value for the 1st parameter (FSC) of the 1st tube was calculated using a histogram matrix comprised of 179 histogram vectors corresponding to the FSC. In the case of redundant parameters such as FSC, SSC, and CD45-ECD, only one parameter with the lowest Λ among them was used.

To validate the result of Λ measurement, the median fluorescence intensities (MFI) of specific parameters with low Λ values were compared, and *t* test was conducted to examine if the differences between AML and non-AML are significant. In addition, t-Distributed Stochastic Neighbor Embedding (t-SNE) on the data matrix comprised of the histogram vectors of a parameter was performed to qualitatively examine the AML discrimination ability of the specific parameters.

#### Marker subset selection

The best subset of parameters was selected via stepwise–forward selection, which is a procedure that begins with an empty dataset and adds parameters to the regression or classification model one by one [17]. In each forward step, the best parameter with the lowest Λ value among the remaining parameters was sequentially added to the dataset, and then the prediction performance of the AML classification model was compared to select the best subset.

In each step of stepwise–forward selection, the classification model was developed using the random-forest algorithm (R package), an ensemble learning method taking multiple decision trees [18]. *F*-measure, which was used in FlowCAP-II challenge, was calculated in order to compare model performance with those of other algorithms proposed in the challenge under the same criteria. The area under the receiver operating characteristic curve, sensitivity, specificity, positive predictive value, and negative predictive value were computed as well.

From the random-forest model, MDG, which is a measure of variable importance for classifying a target, can be calculated. MDG is the basis of Gini impurity, which is the probability of incorrectly classifying a randomly chosen element in a dataset if it were randomly labeled according to the class distribution in the dataset [19]. A decision tree is built by forming each node that leads to the greatest reduction in Gini impurity. Therefore, a higher MDG indicates higher variable importance.

We used MDG to identify specific histogram bin range of each marker of the best subset, which plays an important role in developing an AML classification model. The MDG for 32 bins of each important marker was computed, and then a specific bin range with the highest MDG was identified from each marker histogram. The influence of the specific bins on AML discrimination was examined through heatmap, a graphical representation useful for cross-examining multivariate data.

## Results and discussion

### Feature extraction

The data of each parameter of each FCS file were represented using a histogram vector comprised of 32 bins of equal size. To measure the Λ value of each parameter, a histogram matrix comprised of *n* histogram vectors, which represent each parameter of the samples, was used. In the model development, histogram vectors of two or more parameters of a sample were concatenated into a single large histogram vector. For example, two histogram vectors of FSC and SSC of the 1st sample were combined into a vector with a length of 2*32 (the number of parameters times the number of bins). In the case of the training set for 179 samples, dataset comprised of 178 (rows) by *n*\*32 was used to build up the AML classification model.

### Important marker selection

Λ values calculated from the training data are shown in Table 1. The results suggest that SSC and FSC, which contribute to identification of blasts, have the best ability in AML discrimination. Next, myeloid markers (e.g., CD15, CD34, CD117, and HLA-DR) and lymphocyte markers (e.g., CD38, CD5, CD7, and CD10) were listed in sequential order. The IgG1 and parameter related to the unstained tube showed relatively high Λ values compared to the others.

**Table 2.** Wilks’ lambda of each parameter.

| Rank | Parameter | Wilks' Lambda |
| --- | --- | --- |
| 1 | SSC | 0.2324 |
| 2 | FSC | 0.3263 |
| 3 | CD15-FITC | 0.3442 |
| 4 | CD34-PC5 | 0.3460 |
| 5 | CD117-PE | 0.3667 |
| 6 | HLA DR-FITC | 0.4125 |
| 7 | CD16-PC5 | 0.4368 |
| 8 | CD38-PC7 | 0.4657 |
| 9 | CD56-PC7 | 0.5203 |
| 10 | CD14-FITC | 0.5233 |
| 11 | CD33-PC7 | 0.5349 |
| 12 | CD64-PC5 | 0.5479 |
| 13 | CD5-FITC | 0.5676 |
| 14 | Kappa-FITC | 0.5782 |
| 15 | Lambda-PE | 0.5816 |
| 16 | CD7-FITC | 0.5838 |
| 17 | CD45-ECD | 0.5936 |
| 18 | CD13-PE | 0.6125 |
| 19 | IgG1-FITC | 0.6270 |
| 20 | CD11c-PE | 0.6283 |
| 21 | 7AAD | 0.6297 |
| 22 | CD10-PC7 | 0.6392 |
| 23 | CD3-PC5 | 0.6445 |
| 24 | IgG1-PE | 0.6516 |
| 25 | CD19-PC5 | 0.6537 |
| 26 | CD20-PC7 | 0.6646 |
| 27 | CD8-PC5 | 0.6647 |
| 28 | CD4-PE | 0.6799 |
| 29 | CD19-PE | 0.6920 |
| 30 | FL1 | 0.6942 |
| 31 | FL2 | 0.7023 |
| 32 | CD2-PC7 | 0.7065 |
| 33 | IgG1-PC5 | 0.752 |
| 34 | FL4 | 0.7774 |
| 35 | FL5 | 0.7947 |
| 36 | IgG1-PC7 | 0.8031 |

The order of importance of each parameter based on Λ value supports the general gating strategy which has been used for blast identification, blast lineage assignment, and identification of aberrant immunophenotypic features. In the general gating strategy, the blasts are first gated, and then abnormal myeloid and lymphoid blasts are identified using related lineage-specific markers [6]. The gating of blasts is an important procedure in AML diagnosis because the diagnosis of AML depends on the percent of blasts (>20%) in the bone marrow or blood [20]. For gating of blasts, CD45 versus SSC gating is commonly used because it provides distinct discrimination between the various cell lineages such as lymphocytes, hematogones, monocytes, granulocytes, and myeloid blasts [21].

In this study, CD45 with intermediate Λ value (0.5936) was identified as a less important marker compared to SSC and FSC used for blasts gating. Its Λ value indicates that CD45 does not contribute much to AML discrimination when used alone. Fig 1 shows the MFI of SSC, FSC, and CD45 between non-AML versus AML. MFIs of all parameters and p-value calculated using *t* test are listed in S1 Table. According to the results of the *t* test, there were significant differences in SSC (p-value: 1.66E-10) and FSC (p-value: 2.75E-05) unlike in CD45 (p-value: 0.01). Fig 2 shows the t-SNE results for SSC, FSC, and CD45. In the 2D t-SNE maps for SSC and FSC, AML cluster is relatively well discriminated from non-AML cluster, compared to in the clustering with CD45.

**Fig 1.**
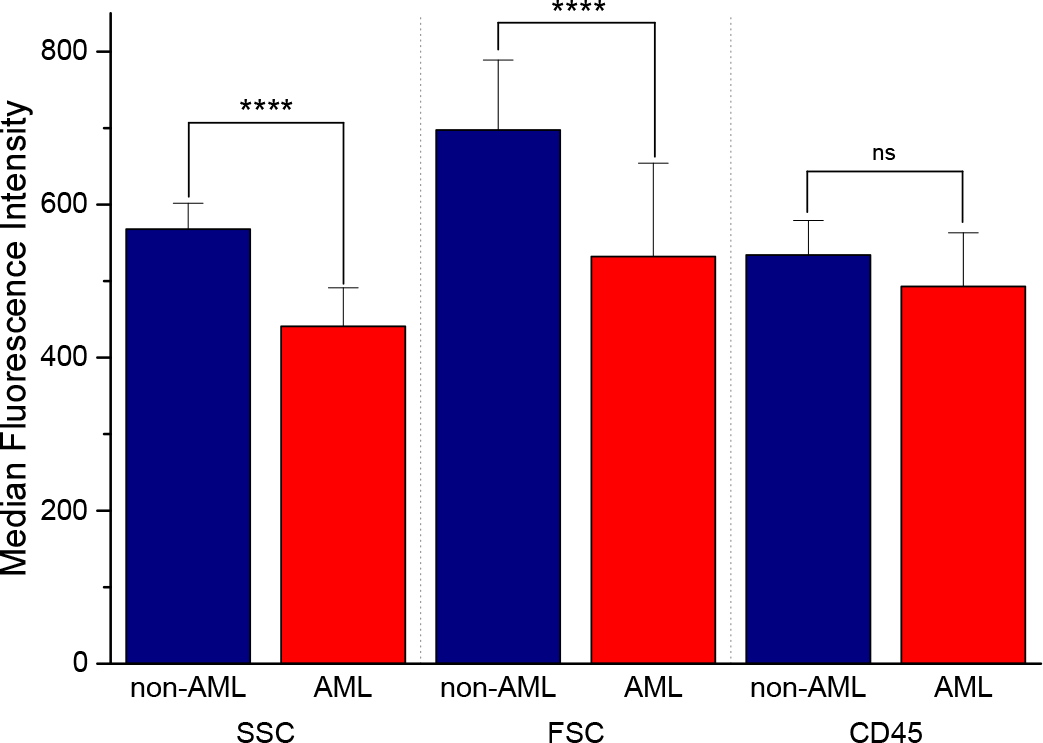
MFI of SSC, FCS, and CD45. ****: p-value < 0.0001, ns: not significant.

**Fig 2.**
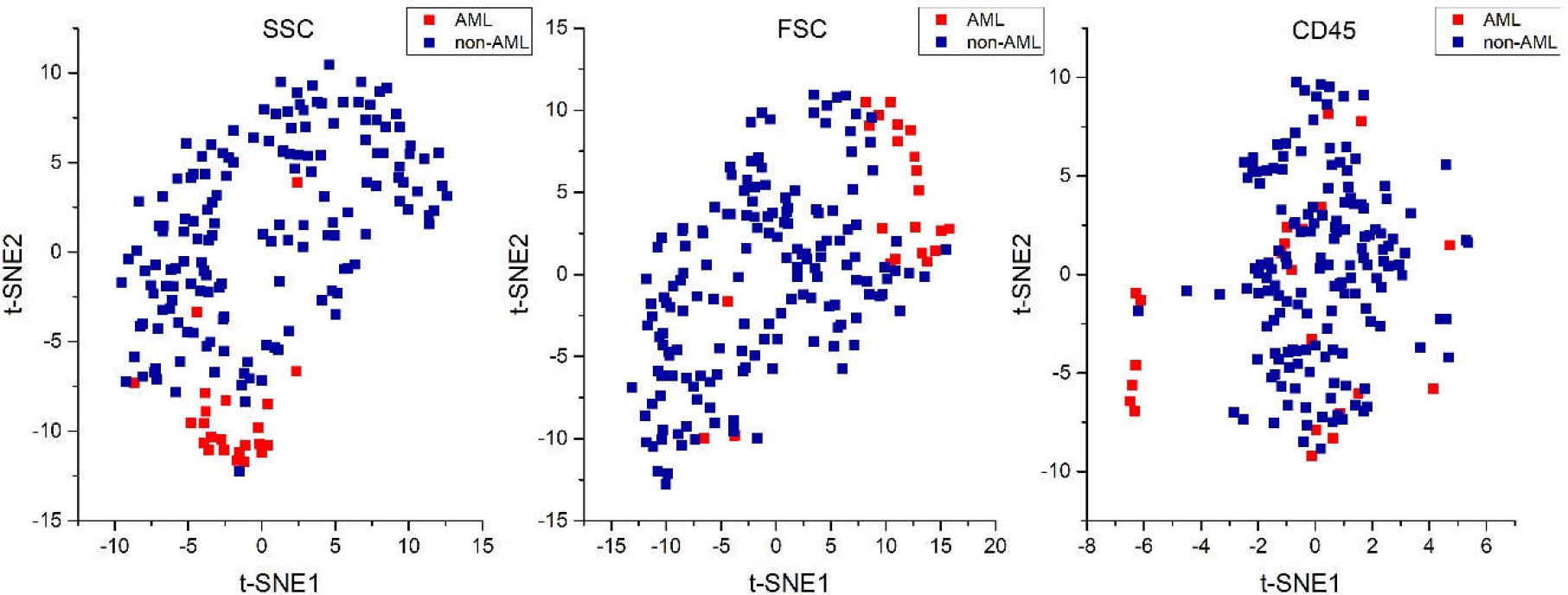
2D t-SNE map for SSC, FCS, and CD45.

Because the Λ value is estimated using only 1D histogram vector of each parameter, the associations between pairs of parameters are not reflected in the Λ calculation. Therefore, further studies on feature extraction to compliment this limitation of Λ-based approach are required. The AML classification challenge of FlowCAP II is not a complex problem because samples from healthy people were used as a control group. To investigate the limitations of the presented approach, a comparison between AML and non-AML that is difficult to distinguish from AML is needed.

### Marker subset selection

In each step of the stepwise–forward selection, the parameter with the smallest Λ value among the parameters was sequentially added to the dataset, and then the AML classification model was developed. Fig 3 shows the model performance (*F*-measure) in each step for the training and testing sets. The parameter sequentially added in each step to the dataset for model development is listed on the y-axis in Fig 3. The numerical performance of the model on the training and testing phases are listed in S2 and S3 Tables, respectively.

**Fig 3.**
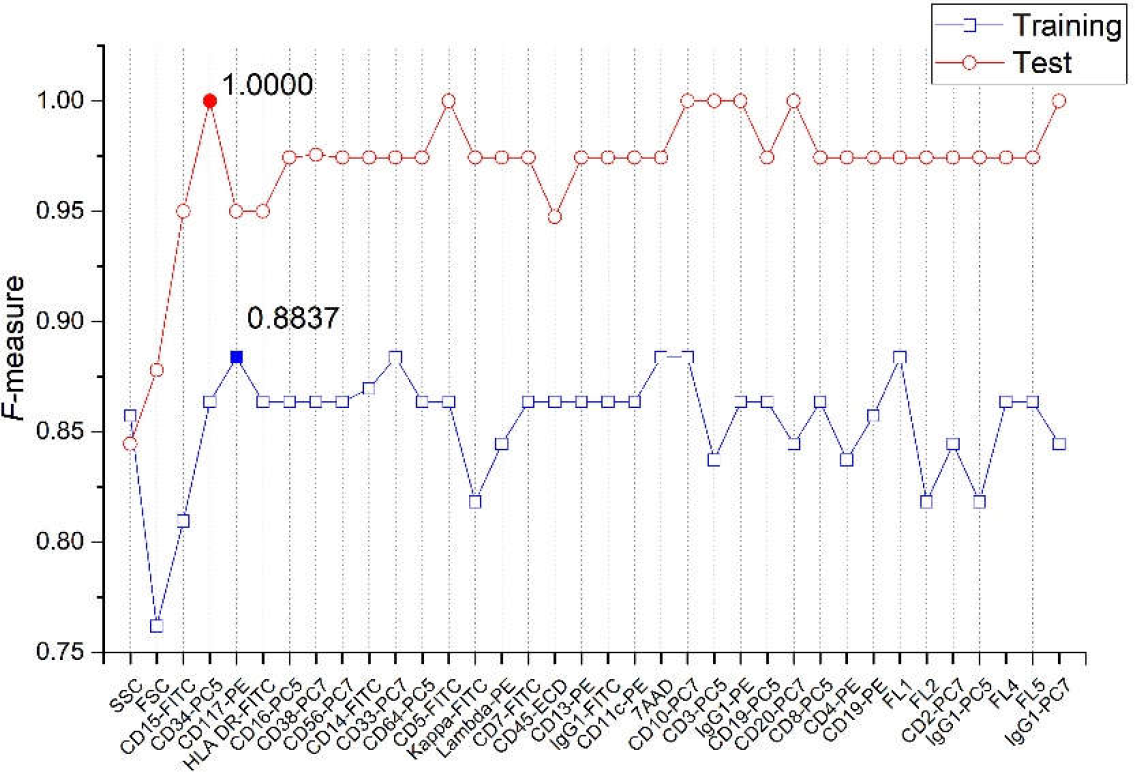
Model performance in each step of stepwise–forward selection.

The AML classification model developed using 5 parameters SSC, FSC, CD15, CD34, and CD117 showed the best performance in the training. In the case of testing, the model correctly predicted all labels of the test set when 4 parameters SSC, FSC, CD15, and CD34 were added to a dataset for model training. CD15, CD34, and CD117 have been used for identification of immature myeloid cells [6]. It indicates that the parameters associated with blasts gating and immature myeloid cells play an important role in discriminating AML and non-AML.

In both training and testing, the model performance was not significantly improved after 7 or more parameters were used. In particular, the model performance spiked when SSC, FSC, CD15, and CD34 were sequentially added to the modelling dataset in stepwise–forward selection. This means that a few markers significantly contribute to discriminate AML, and a combination of these markers is more effective than an individual marker. In the case of the training results, the model performance tends to decrease after markers related to the identification of cellular subsets of the human immune system and fluorescence channels, corresponding to unstained tube#8, were appended to the modelling dataset. The addition of such markers, which were less important compared to the markers related to the immaturity of myeloid cells in Wilks’ lambda analysis, hindered model performance.

Although the blasts gating-related markers were mainly identified as important descriptors for AML classification in this study, the additional markers must be considered for the blast lineage assignment and identification of aberrant immunophenotypic features in the clinical testing.

In all step of the stepwise forward selection, model performance difference between training and test dataset was observed. The discrepancy between training and test results affects the generalization performance of the model. We performed t-SNE on the dataset comprised of the best subset in order to find out cause leading to the discrepancy. S1 Fig shows the 2-D t-SNE result. In the case of training, four AML samples (sample no. 5, 7, 101, and 116), which were frequently misclassified as non-AML class in model training, were in domain of non-AML training dataset. These AML samples led to an increase in the false negative rate. By contrast, AML and non-AML samples of the test dataset were clearly distributed within the domain of each corresponding group formed by the training set. Because of this, the model exactly classified AML and non-AML of test set. In the test set, no. 340 sample is located at border between training and test data domain. This sample, which may have a preleukemic condition rather than AML, was frequently misclassified by other algorithms in the FlowCAP-II challenge. Our t-SNE result reveals that there are samples (no. 5, 7, 101, and 116) which should be examined in detail and the random data split in the FlowCAP-II challenge influenced the performance.

As shown in S2 Fig, the proposed model showed a relatively good performance among the other models used in FlowCAP-II AML classification challenge.

Fig 4 shows a color-coded heatmap for model training dataset-comprised histograms of the identified 4 important markers. It consists of 179 rows, with each row corresponding to each sample, and 128 columns related to the histogram of each marker. AML and non-AML groups are separated by a horizontal white border. Blue, white, and red colors indicate low, intermediate, and high frequencies, respectively, wherein frequency is the number of cells (or events) falling within each bin of the histogram. Four colors at the top of the heat map indicate each related important parameter.

**Fig 4.**
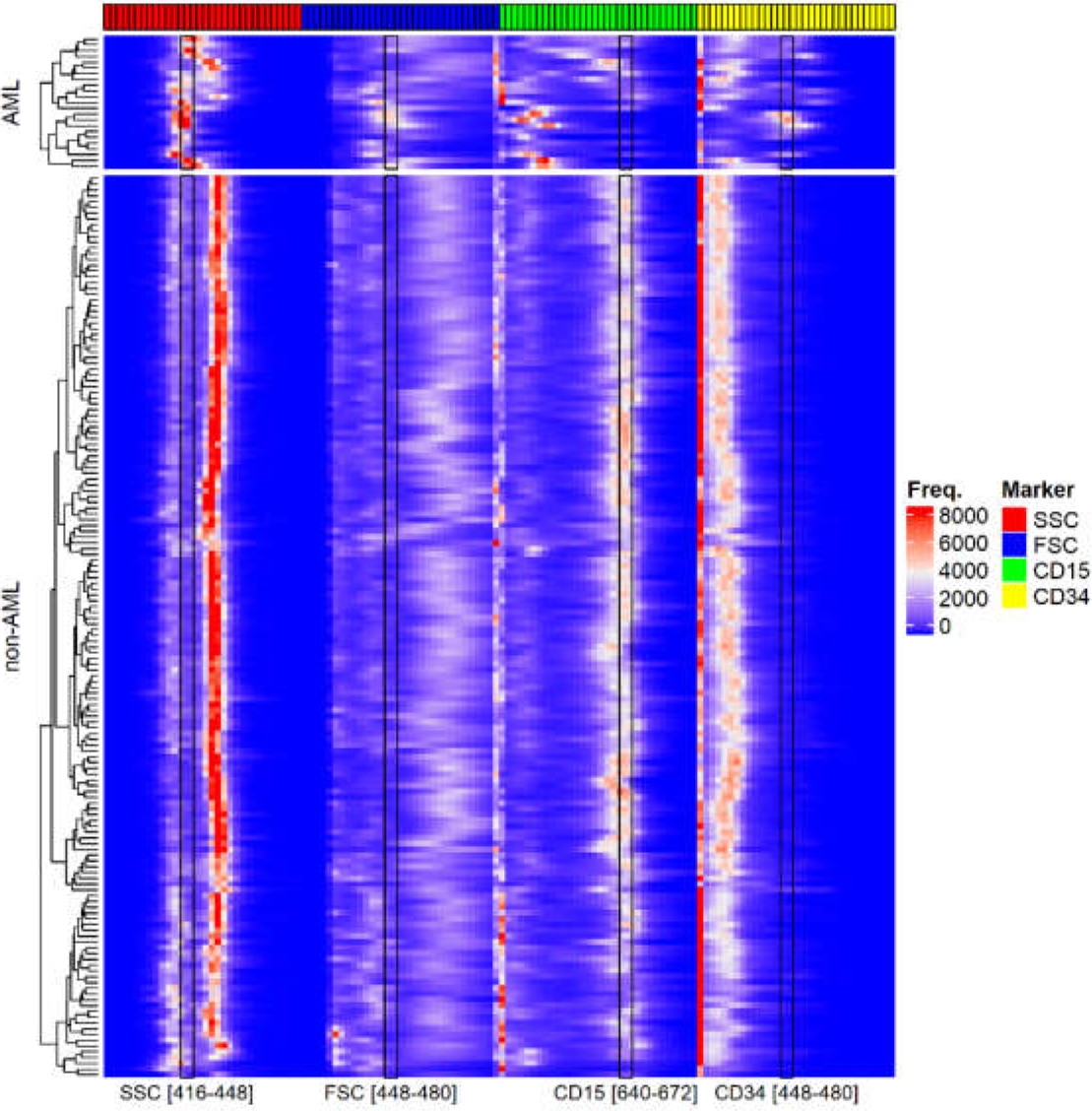
Heatmap of input matrix for the best subset.

The calculated MDGs are listed in S4 Table. MDG results show that each bin has a different AML discrimination ability. From the 4 markers, 4 important bin ranges were respectively identified: SSC [416-448], FSC [448-480], CD15 [640-672], and CD34 [448-480]. The rectangles on the heatmap show the identified bin ranges with the high MDS of each marker. As shown in Fig 4, the ranges display different colors between AML and non-AML. In the case of SSC and FSC, the cell frequency (or expression) of the identified range in AML is higher than in non-AML because the percentage of blasts increases in AML. Furthermore, the population of the blast cells is expanded between low and intermediate levels of both SSC and FSC because of the low granularity and small to intermediate sizes of the blast cells [22]. For CD15, the expression in the range [640-672] in AML was relatively lower than in non-AML. CD15 shows a generally negative expression in M0 and M1 AML [23]. CD34 is expressed on hematopoietic stem cells, endothelial cells, leukemic blast cells, and normal hematopoietic progenitors [3]. The number of CD34^+^ cells reflects the number of immature myeloid and erythroid cells, in particular the myeloid blast, released into the peripheral blood [24]. Therefore, patients with blasts exhibit higher expression of CD34 than do patients without blasts. In the heatmap, CD34 exhibits relatively higher expression in the identified range [448-480] than in non-AML.

The drawbacks of manual gating have motivated the development of automated analysis methods for cell-population identification, sample classification, and rare cell identification [13,25,26]. In terms of sample classification, there have been efforts to build up sample-classification models [27–30].

A pipeline for model development typically comprises of feature extraction from each sample, important-feature (parameter) identification, model development, and interpretation. For the feature extraction from each specimen, clustering algorithms or expression of each fluorescence channel is used. In the case of the clustering algorithms, *n* clusters in each sample are identified and then feature values (e.g., a fraction of events, MFI, and mean FSC and SSC values) of each identified cluster are extracted, while another method uses expression profiling information (e.g., percentage of positive cells for each antigen, percentage of blast cells within the CD45/SSC gate, and positive or negative expression) as feature values. Important-parameter identification is an essential processing step in the pipeline because it reduces the complexity of the model and leads to better interpretations. The important parameters are identified using stepwise linear discriminant analysis based on least absolute shrinkage and selection operator (LASSO) and Wilks’ lambda. In particular, both important-parameter selection and regularization of the linear-regression model are simultaneously performed in the discriminant analysis. Finally, interpretation is carried out using statistics of identified parameters.

The aforementioned feature-extraction methods only use limited representative features of each identified cluster or each fluorescence signal expression for antigen. Although thousands of events in each channel are generally divided into positive and negative by an experimentally determined threshold, there is still informative event expression change over the range of each channel for a specific antibody. From such information, complementary and alternative features can be produced. Because of this, histogram, which displays a single measurement of flow-cytometry parameter, was used in this study. Those features could complement each other. One of the best merits of the histogram is that it allows exploration of expression change over a specific range. Identifying the specific ranges can contribute not only to elaborating the classification model as well as the decision-support system and provide but also to helping establish precise diagnostic criteria. We used MDG criteria based on Gini impurity to spot the specific ranges. As a result, it was confirmed that each range has a different discriminate ability and that particular ranges with high MDG significantly influence AML classification in the model.

## Conclusions

We estimated the importance of each marker via Wilks’ lambda and then identified the best subset of markers using stepwise–forward selection for AML classification model development. As a result, markers, which are associated with the blasts gating and identification of immature myeloid cells, were identified to contribute significantly to AML classification, and the classification model developed using their combination showed good predictive power. In addition, we highlighted decision-support range of the fluorescent signal for the identified important parameters, which significantly contribute to AML classification, through MDG supported in random forest. Because the methodology presented in this study can not only estimate the importance of each marker but can also identify the best subset of markers and the specific ranges, we expect that it would contribute to *in silico* modeling using flow- and mass-cytometry readout as well as panel design for sample classification.

Samples obtained from healthy people were used as a control group in this study. To validate the limitation and usefulness of the presented methodology in terms of sample classification, additional study on a dataset with samples, which are difficult to distinguish clinically from AML samples, should be performed. In addition, because the Λ value is estimated using only a 1D histogram vector of each parameter, the association between pairs of parameters is not reflected in the Λ results. To overcome this problem, a more improved feature-extraction method is required.

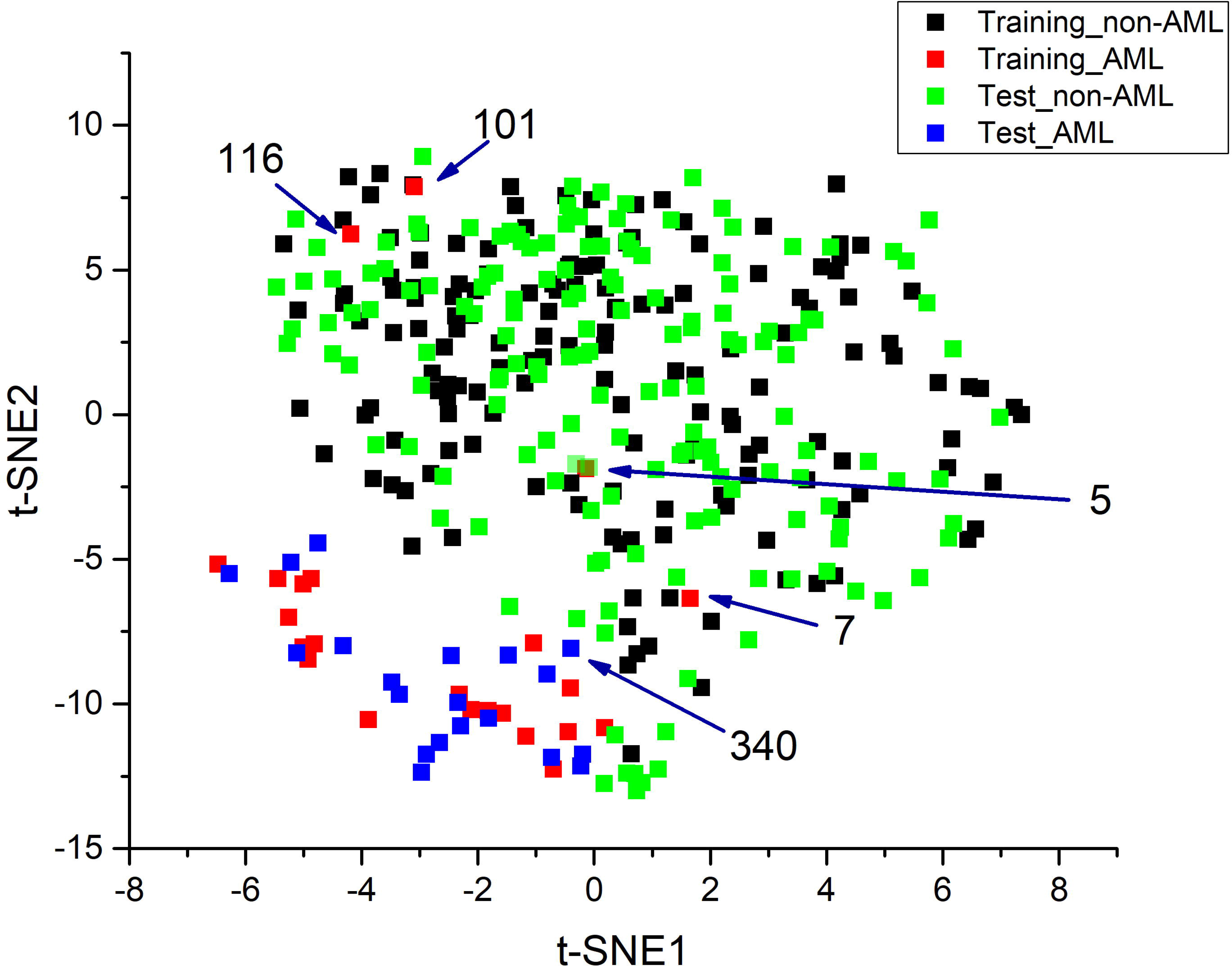

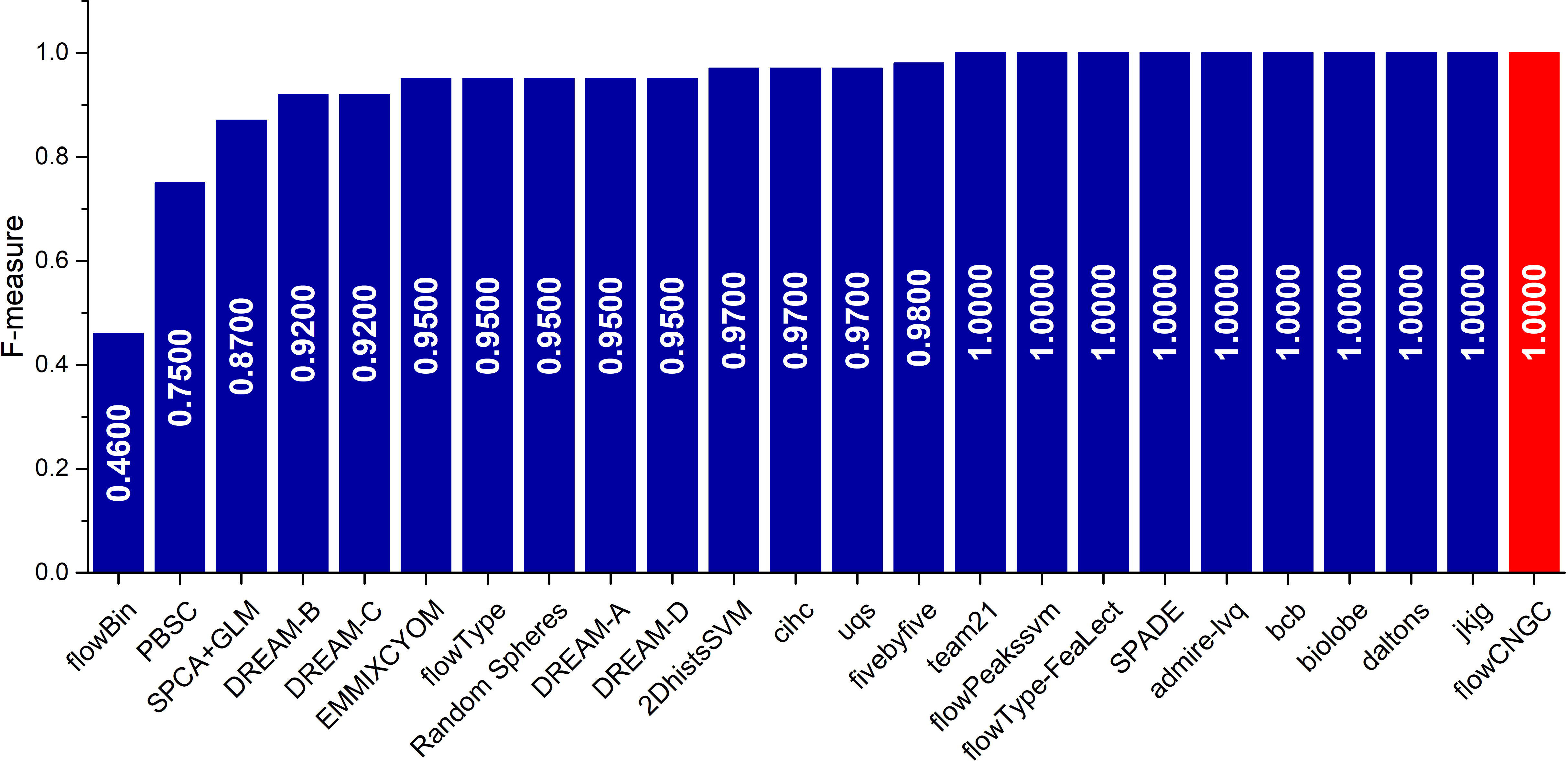

## Acknowledgements

The authors appreciate CNGC members and other collaborators for their contributions.

## Supplementary data

Supporting information.docx: This file provides supplementary tables and figures.

