## Supporting information for "Marker subset selection and decision support range identification for acute myeloid leukemia classification model development with multiparameter flow cytometry"

Supporting information includes:
S1 Table. MFI of all parameters and *p*-value calculated using *t* test

S2 Table. Model performance for training set in each step of stepwise forward selection

S3 Table. Model performance for test set in each step of stepwise forward selection

S4 Table. Important marker bin range and mean decrease in Gini

S1 Fig. 2D t-SNE plot for training and test set

S2 Fig. Model performance comparison with algorithms used in FlowCAP-II Challenge

S1 Table. MFI of all parameter and *p*-value calculated using *t* test

| **No.** | **Parameter** | **AML** | | **non-AML** | | ***p*-value** |
| --- | --- | --- | --- | --- | --- | --- |
|  |  | **Median** | **SD** | **Median** | **SD** |  |
| 1 | SSC | 441.0000 | 50.3817 | 568.0000 | 33.6549 | 1.66E-10 |
| 2 | FSC | 532.0000 | 122.0409 | 697.5000 | 91.4463 | 2.75E-05 |
| 3 | CD15-FITC | 251.0000 | 147.1892 | 584.5000 | 98.3116 | 6.63E-09 |
| 4 | CD34-PC5 | 210.0000 | 143.3781 | 116.0000 | 34.7504 | 0.0009 |
| 5 | CD117-PE | 276.0000 | 78.2391 | 208.5000 | 39.4915 | 6.17E-05 |
| 6 | HLA DR-FITC | 350.0000 | 149.8541 | 251.5000 | 47.2385 | 0.0026 |
| 7 | CD16-PC5 | 88.0000 | 120.2692 | 357.5000 | 164.1421 | 2.88E-11 |
| 8 | CD38-PC7 | 386.0000 | 105.4701 | 273.5000 | 54.1296 | 0.0002 |
| 9 | CD56-PC7 | 63.0000 | 121.6648 | 12.5000 | 27.9291 | 0.0038 |
| 10 | CD14-FITC | 267.0000 | 103.7197 | 305.5000 | 57.4106 | 0.0765 |
| 11 | CD33-PC7 | 357.0000 | 115.3863 | 313.5000 | 73.8985 | 0.1083 |
| 12 | CD64-PC5 | 279.0000 | 161.0872 | 257.0000 | 89.7994 | 0.9218 |
| 13 | CD5-FITC | 219.0000 | 71.8018 | 247.0000 | 41.8168 | 0.1373 |
| 14 | Kappa-FITC | 353.0000 | 103.2710 | 368.0000 | 59.2987 | 0.0466 |
| 15 | Lambda-PE | 377.0000 | 119.7949 | 392.0000 | 64.1720 | 0.1550 |
| 16 | CD7-FITC | 357.0000 | 105.9434 | 322.5000 | 51.8503 | 0.1918 |
| 17 | CD45-ECD | 493.0000 | 70.2663 | 534.0000 | 45.2695 | 0.0100 |
| 18 | CD13-PE | 421.0000 | 133.7059 | 385.5000 | 87.1620 | 0.7739 |
| 19 | IgG1-FITC | 192.0000 | 57.3249 | 235.0000 | 47.8723 | 0.0046 |
| 20 | CD11c-PE | 274.5000 | 104.1956 | 307.5000 | 94.1832 | 0.1616 |
| 21 | 7AAD | 157.0000 | 29.0160 | 191.0000 | 48.4896 | 3.71E-05 |
| 22 | CD10-PC7 | 47.0000 | 46.1336 | 94.5000 | 60.2379 | 9.66E-06 |
| 23 | CD3-PC5 | 114.0000 | 53.3837 | 140.0000 | 34.1454 | 0.019 |
| 24 | IgG1-PE | 173.0000 | 45.2703 | 196.5000 | 37.1716 | 0.020 |
| 25 | CD19-PC5 | 95.0000 | 76.9307 | 95.0000 | 44.4103 | 0.793 |
| 26 | CD20-PC7 | 100.0000 | 50.1251 | 106.0000 | 37.8641 | 0.862 |
| 27 | CD8-PC5 | 95.0000 | 50.9487 | 110.5000 | 33.3227 | 0.029 |
| 28 | CD4-PE | 200.0000 | 61.5406 | 205.0000 | 35.2502 | 0.807 |
| 29 | CD19-PE | 177.0000 | 62.0871 | 206.0000 | 35.5158 | 0.112 |
| 30 | FL1 | 192.0000 | 54.4815 | 236.0000 | 47.2041 | 0.003 |
| 31 | FL2 | 166.0000 | 40.3988 | 204.0000 | 34.1372 | 0.001 |
| 32 | CD2-PC7 | 75.0000 | 32.8713 | 69.0000 | 24.7226 | 0.592 |
| 33 | IgG1-PC5 | 99.0000 | 51.5755 | 119.0000 | 34.6178 | 0.017 |
| 34 | FL4 | 76.0000 | 40.8976 | 107.0000 | 27.6755 | 0.064 |
| 35 | FL5 | 44.0000 | 21.5342 | 57.0000 | 23.2155 | 0.061 |
| 36 | IgG1-PC7 | 54.0000 | 27.6072 | 64.5000 | 28.9562 | 0.143 |

S2 Table. Model performance for training set in each step of stepwise forward selection

| **Step** | **Parameter Added** | **TP** | **FP** | **FN** | **TN** | **Sensitivity** | **Specificity** | **PPV** | **NPV** | **F-measure** | **Balanced Accuracy** |
| --- | --- | --- | --- | --- | --- | --- | --- | --- | --- | --- | --- |
| 1 | SSC | 18 | 1 | 5 | 155 | 0.7826 | 0.9936 | 0.9474 | 0.9688 | 0.8571 | 0.8881 |
| 2 | FSC | 16 | 3 | 7 | 153 | 0.6957 | 0.9808 | 0.8421 | 0.9563 | 0.7619 | 0.8382 |
| 3 | CD15-FITC | 17 | 2 | 6 | 154 | 0.7391 | 0.9872 | 0.8947 | 0.9625 | 0.8095 | 0.8632 |
| 4 | CD34-PC5 | 19 | 2 | 4 | 154 | 0.8261 | 0.9872 | 0.9048 | 0.9747 | 0.8636 | 0.9066 |
| 5 | CD117-PE | 19 | 1 | 4 | 155 | 0.8261 | 0.9936 | 0.9500 | 0.9748 | 0.8837 | 0.9098 |
| 6 | HLA DR-FITC | 19 | 2 | 4 | 154 | 0.8261 | 0.9872 | 0.9048 | 0.9747 | 0.8636 | 0.9066 |
| 7 | CD16-PC5 | 19 | 2 | 4 | 154 | 0.8261 | 0.9872 | 0.9048 | 0.9747 | 0.8636 | 0.9066 |
| 8 | CD38-PC7 | 19 | 2 | 4 | 154 | 0.8261 | 0.9872 | 0.9048 | 0.9747 | 0.8636 | 0.9066 |
| 9 | CD56-PC7 | 19 | 2 | 4 | 154 | 0.8261 | 0.9872 | 0.9048 | 0.9747 | 0.8636 | 0.9066 |
| 10 | CD14-FITC | 20 | 3 | 3 | 153 | 0.8696 | 0.9808 | 0.8696 | 0.9808 | 0.8696 | 0.9252 |
| 11 | CD33-PC7 | 19 | 1 | 4 | 155 | 0.8261 | 0.9936 | 0.9500 | 0.9748 | 0.8837 | 0.9098 |
| 12 | CD64-PC5 | 19 | 2 | 4 | 154 | 0.8261 | 0.9872 | 0.9048 | 0.9747 | 0.8636 | 0.9066 |
| 13 | CD5-FITC | 19 | 2 | 4 | 154 | 0.8261 | 0.9872 | 0.9048 | 0.9747 | 0.8636 | 0.9066 |
| 14 | Kappa-FITC | 18 | 3 | 5 | 153 | 0.7826 | 0.9808 | 0.8571 | 0.9684 | 0.8182 | 0.8817 |
| 15 | Lambda-PE | 19 | 3 | 4 | 153 | 0.8261 | 0.9808 | 0.8636 | 0.9745 | 0.8444 | 0.9034 |
| 16 | CD7-FITC | 19 | 2 | 4 | 154 | 0.8261 | 0.9872 | 0.9048 | 0.9747 | 0.8636 | 0.9066 |
| 17 | CD45-ECD | 19 | 2 | 4 | 154 | 0.8261 | 0.9872 | 0.9048 | 0.9747 | 0.8636 | 0.9066 |
| 18 | CD13-PE | 19 | 2 | 4 | 154 | 0.8261 | 0.9872 | 0.9048 | 0.9747 | 0.8636 | 0.9066 |
| 19 | IgG1-FITC | 19 | 2 | 4 | 154 | 0.8261 | 0.9872 | 0.9048 | 0.9747 | 0.8636 | 0.9066 |
| 20 | CD11c-PE | 19 | 2 | 4 | 154 | 0.8261 | 0.9872 | 0.9048 | 0.9747 | 0.8636 | 0.9066 |
| 21 | 7AAD | 19 | 1 | 4 | 155 | 0.8261 | 0.9936 | 0.9500 | 0.9748 | 0.8837 | 0.9098 |
| 22 | CD10-PC7 | 19 | 1 | 4 | 155 | 0.8261 | 0.9936 | 0.9500 | 0.9748 | 0.8837 | 0.9098 |
| 23 | CD3-PC5 | 18 | 2 | 5 | 154 | 0.7826 | 0.9872 | 0.9000 | 0.9686 | 0.8372 | 0.8849 |
| 24 | IgG1-PE | 19 | 2 | 4 | 154 | 0.8261 | 0.9872 | 0.9048 | 0.9747 | 0.8636 | 0.9066 |
| 25 | CD19-PC5 | 19 | 2 | 4 | 154 | 0.8261 | 0.9872 | 0.9048 | 0.9747 | 0.8636 | 0.9066 |
| 26 | CD20-PC7 | 19 | 3 | 4 | 153 | 0.8261 | 0.9808 | 0.8636 | 0.9745 | 0.8444 | 0.9034 |
| 27 | CD8-PC5 | 19 | 2 | 4 | 154 | 0.8261 | 0.9872 | 0.9048 | 0.9747 | 0.8636 | 0.9066 |
| 28 | CD4-PE | 18 | 2 | 5 | 154 | 0.7826 | 0.9872 | 0.9000 | 0.9686 | 0.8372 | 0.8849 |
| 29 | CD19-PE | 18 | 1 | 5 | 155 | 0.7826 | 0.9936 | 0.9474 | 0.9688 | 0.8571 | 0.8881 |
| 30 | FL1 | 19 | 1 | 4 | 155 | 0.8261 | 0.9936 | 0.9500 | 0.9748 | 0.8837 | 0.9098 |
| 31 | FL2 | 18 | 3 | 5 | 153 | 0.7826 | 0.9808 | 0.8571 | 0.9684 | 0.8182 | 0.8817 |
| 32 | CD2-PC7 | 19 | 3 | 4 | 153 | 0.8261 | 0.9808 | 0.8636 | 0.9745 | 0.8444 | 0.9034 |
| 33 | IgG1-PC5 | 18 | 3 | 5 | 153 | 0.7826 | 0.9808 | 0.8571 | 0.9684 | 0.8182 | 0.8817 |
| 34 | FL4 | 19 | 2 | 4 | 154 | 0.8261 | 0.9872 | 0.9048 | 0.9747 | 0.8636 | 0.9066 |
| 35 | FL5 | 19 | 2 | 4 | 154 | 0.8261 | 0.9872 | 0.9048 | 0.9747 | 0.8636 | 0.9066 |
| 36 | IgG1-PC7 | 19 | 3 | 4 | 153 | 0.8261 | 0.9808 | 0.8636 | 0.9745 | 0.8444 | 0.9034 |

S3 Table. Model performance for test set in each step of stepwise forward selection

| **Step** | **Parameter Added** | **TP** | **FP** | **FN** | **TN** | **Sensitivity** | **Specificity** | **PPV** | **NPV** | **F-measure** | **Balanced Accuracy** |
| --- | --- | --- | --- | --- | --- | --- | --- | --- | --- | --- | --- |
| 1 | SSC | 19 | 6 | 1 | 154 | 0.9500 | 0.9625 | 0.7600 | 0.9935 | 0.8444 | 0.9563 |
| 2 | FSC | 18 | 3 | 2 | 157 | 0.9000 | 0.9813 | 0.8571 | 0.9874 | 0.8780 | 0.9406 |
| 3 | CD15-FITC | 19 | 1 | 1 | 159 | 0.9500 | 0.9938 | 0.9500 | 0.9938 | 0.9500 | 0.9719 |
| 4 | CD34-PC5 | 20 | 0 | 0 | 160 | 1.0000 | 1.0000 | 1.0000 | 1.0000 | 1.0000 | 1.0000 |
| 5 | CD117-PE | 19 | 1 | 1 | 159 | 0.9500 | 0.9938 | 0.9500 | 0.9938 | 0.9500 | 0.9719 |
| 6 | HLA DR-FITC | 19 | 1 | 1 | 159 | 0.9500 | 0.9938 | 0.9500 | 0.9938 | 0.9500 | 0.9719 |
| 7 | CD16-PC5 | 19 | 0 | 1 | 160 | 0.9500 | 1.0000 | 1.0000 | 0.9938 | 0.9744 | 0.9750 |
| 8 | CD38-PC7 | 20 | 1 | 0 | 159 | 1.0000 | 0.9938 | 0.9524 | 1.0000 | 0.9756 | 0.9969 |
| 9 | CD56-PC7 | 19 | 0 | 1 | 160 | 0.9500 | 1.0000 | 1.0000 | 0.9938 | 0.9744 | 0.9750 |
| 10 | CD14-FITC | 19 | 0 | 1 | 160 | 0.9500 | 1.0000 | 1.0000 | 0.9938 | 0.9744 | 0.9750 |
| 11 | CD33-PC7 | 19 | 0 | 1 | 160 | 0.9500 | 1.0000 | 1.0000 | 0.9938 | 0.9744 | 0.9750 |
| 12 | CD64-PC5 | 19 | 0 | 1 | 160 | 0.9500 | 1.0000 | 1.0000 | 0.9938 | 0.9744 | 0.9750 |
| 13 | CD5-FITC | 20 | 0 | 0 | 160 | 1.0000 | 1.0000 | 1.0000 | 1.0000 | 1.0000 | 1.0000 |
| 14 | Kappa-FITC | 19 | 0 | 1 | 160 | 0.9500 | 1.0000 | 1.0000 | 0.9938 | 0.9744 | 0.9750 |
| 15 | Lambda-PE | 19 | 0 | 1 | 160 | 0.9500 | 1.0000 | 1.0000 | 0.9938 | 0.9744 | 0.9750 |
| 16 | CD7-FITC | 19 | 0 | 1 | 160 | 0.9500 | 1.0000 | 1.0000 | 0.9938 | 0.9744 | 0.9750 |
| 17 | CD45-ECD | 18 | 0 | 2 | 160 | 0.9000 | 1.0000 | 1.0000 | 0.9877 | 0.9474 | 0.9500 |
| 18 | CD13-PE | 19 | 0 | 1 | 160 | 0.9500 | 1.0000 | 1.0000 | 0.9938 | 0.9744 | 0.9750 |
| 19 | IgG1-FITC | 19 | 0 | 1 | 160 | 0.9500 | 1.0000 | 1.0000 | 0.9938 | 0.9744 | 0.9750 |
| 20 | CD11c-PE | 19 | 0 | 1 | 160 | 0.9500 | 1.0000 | 1.0000 | 0.9938 | 0.9744 | 0.9750 |
| 21 | 7AAD | 19 | 0 | 1 | 160 | 0.9500 | 1.0000 | 1.0000 | 0.9938 | 0.9744 | 0.9750 |
| 22 | CD10-PC7 | 20 | 0 | 0 | 160 | 1.0000 | 1.0000 | 1.0000 | 1.0000 | 1.0000 | 1.0000 |
| 23 | CD3-PC5 | 20 | 0 | 0 | 160 | 1.0000 | 1.0000 | 1.0000 | 1.0000 | 1.0000 | 1.0000 |
| 24 | IgG1-PE | 20 | 0 | 0 | 160 | 1.0000 | 1.0000 | 1.0000 | 1.0000 | 1.0000 | 1.0000 |
| 25 | CD19-PC5 | 19 | 0 | 1 | 160 | 0.9500 | 1.0000 | 1.0000 | 0.9938 | 0.9744 | 0.9750 |
| 26 | CD20-PC7 | 20 | 0 | 0 | 160 | 1.0000 | 1.0000 | 1.0000 | 1.0000 | 1.0000 | 1.0000 |
| 27 | CD8-PC5 | 19 | 0 | 1 | 160 | 0.9500 | 1.0000 | 1.0000 | 0.9938 | 0.9744 | 0.9750 |
| 28 | CD4-PE | 19 | 0 | 1 | 160 | 0.9500 | 1.0000 | 1.0000 | 0.9938 | 0.9744 | 0.9750 |
| 29 | CD19-PE | 19 | 0 | 1 | 160 | 0.9500 | 1.0000 | 1.0000 | 0.9938 | 0.9744 | 0.9750 |
| 30 | FL1 | 19 | 0 | 1 | 160 | 0.9500 | 1.0000 | 1.0000 | 0.9938 | 0.9744 | 0.9750 |
| 31 | FL2 | 19 | 0 | 1 | 160 | 0.9500 | 1.0000 | 1.0000 | 0.9938 | 0.9744 | 0.9750 |
| 32 | CD2-PC7 | 19 | 0 | 1 | 160 | 0.9500 | 1.0000 | 1.0000 | 0.9938 | 0.9744 | 0.9750 |
| 33 | IgG1-PC5 | 19 | 0 | 1 | 160 | 0.9500 | 1.0000 | 1.0000 | 0.9938 | 0.9744 | 0.9750 |
| 34 | FL4 | 19 | 0 | 1 | 160 | 0.9500 | 1.0000 | 1.0000 | 0.9938 | 0.9744 | 0.9750 |
| 35 | FL5 | 19 | 0 | 1 | 160 | 0.9500 | 1.0000 | 1.0000 | 0.9938 | 0.9744 | 0.9750 |
| 36 | IgG1-PC7 | 20 | 0 | 0 | 160 | 1.0000 | 1.0000 | 1.0000 | 1.0000 | 1.0000 | 1.0000 |

S4 Table. Important marker bin range and mean decrease in Gini

| **Rank** | **Marker** | **Bin** | **Range** | **MDG** | **Marker** | **Bin** | **Range** | **MDG** | **Marker** | **Bin** | **Range** | **MDG** | **Marker** | **Bin** | **Range** | **MDG** |
| --- | --- | --- | --- | --- | --- | --- | --- | --- | --- | --- | --- | --- | --- | --- | --- | --- |
| 1 | SSC | V14 | 416-448 | 2.8371 | FSC | V47 | 448-480 | 1.2576 | CD15 | V85 | 640-672 | 1.2996 | CD34 | V111 | 448-480 | 1.8108 |
| 2 | SSC | V19 | 576-608 | 2.0301 | FSC | V38 | 160-192 | 0.6703 | CD15 | V86 | 672-704 | 0.6721 | CD34 | V110 | 416-448 | 1.6603 |
| 3 | SSC | V15 | 448-480 | 1.1789 | FSC | V57 | 768-800 | 0.4900 | CD15 | V74 | 288-320 | 0.6242 | CD34 | V115 | 576-608 | 1.5701 |
| 4 | SSC | V20 | 608-640 | 1.0204 | FSC | V39 | 192-224 | 0.4506 | CD15 | V84 | 608-640 | 0.5957 | CD34 | V112 | 480-512 | 1.3793 |
| 5 | SSC | V13 | 384-416 | 0.8061 | FSC | V40 | 224-256 | 0.3666 | CD15 | V71 | 192-224 | 0.4148 | CD34 | V116 | 608-640 | 1.3404 |
| 6 | SSC | V23 | 704-736 | 0.5194 | FSC | V58 | 800-832 | 0.3511 | CD15 | V76 | 352-384 | 0.4022 | CD34 | V114 | 544-576 | 1.2824 |
| 7 | SSC | V22 | 672-704 | 0.4334 | FSC | V59 | 832-864 | 0.3476 | CD15 | V70 | 160-192 | 0.3777 | CD34 | V109 | 384-416 | 1.0415 |
| 8 | SSC | V18 | 544-576 | 0.4129 | FSC | V56 | 736-768 | 0.3214 | CD15 | V78 | 416-448 | 0.3517 | CD34 | V113 | 512-544 | 0.7710 |
| 9 | SSC | V21 | 640-672 | 0.2892 | FSC | V48 | 480-512 | 0.2714 | CD15 | V72 | 224-256 | 0.3451 | CD34 | V108 | 352-384 | 0.5575 |
| 10 | SSC | V24 | 736-768 | 0.2187 | FSC | V41 | 256-288 | 0.2437 | CD15 | V83 | 576-608 | 0.3263 | CD34 | V101 | 128-160 | 0.3605 |
| 11 | SSC | V16 | 480-512 | 0.1797 | FSC | V46 | 416-448 | 0.1755 | CD15 | V80 | 480-512 | 0.3189 | CD34 | V117 | 640-672 | 0.3356 |
| 12 | SSC | V9 | 256-288 | 0.1303 | FSC | V60 | 864-896 | 0.1528 | CD15 | V73 | 256-288 | 0.3130 | CD34 | V99 | 64-96 | 0.3276 |
| 13 | SSC | V11 | 320-352 | 0.0984 | FSC | V62 | 928-960 | 0.1489 | CD15 | V81 | 512-544 | 0.2655 | CD34 | V98 | 32-64 | 0.2821 |
| 14 | SSC | V26 | 800-832 | 0.0885 | FSC | V37 | 128-160 | 0.1416 | CD15 | V82 | 544-576 | 0.2545 | CD34 | V100 | 96-128 | 0.2377 |
| 15 | SSC | V10 | 288-320 | 0.0878 | FSC | V49 | 512-544 | 0.1401 | CD15 | V79 | 448-480 | 0.1997 | CD34 | V102 | 160-192 | 0.2120 |
| 16 | SSC | V32 | 992-1024 | 0.0833 | FSC | V61 | 896-928 | 0.1048 | CD15 | V75 | 320-352 | 0.1863 | CD34 | V118 | 672-704 | 0.1845 |
| 17 | SSC | V7 | 192-224 | 0.0769 | FSC | V55 | 704-736 | 0.0951 | CD15 | V66 | 32-64 | 0.1452 | CD34 | V119 | 704-736 | 0.1749 |
| 18 | SSC | V8 | 224-256 | 0.0741 | FSC | V50 | 544-576 | 0.0903 | CD15 | V67 | 64-96 | 0.1254 | CD34 | V103 | 192-224 | 0.1539 |
| 19 | SSC | V25 | 768-800 | 0.0740 | FSC | V42 | 288-320 | 0.0865 | CD15 | V89 | 768-800 | 0.0940 | CD34 | V97 | 0-32 | 0.1532 |
| 20 | SSC | V12 | 352-384 | 0.0575 | FSC | V64 | 992-1024 | 0.0837 | CD15 | V88 | 736-768 | 0.0936 | CD34 | V107 | 320-352 | 0.1216 |
| 21 | SSC | V30 | 928-960 | 0.0571 | FSC | V44 | 352-384 | 0.0785 | CD15 | V77 | 384-416 | 0.0898 | CD34 | V120 | 736-768 | 0.0680 |
| 22 | SSC | V29 | 896-928 | 0.0559 | FSC | V51 | 576-608 | 0.0748 | CD15 | V68 | 96-128 | 0.0810 | CD34 | V105 | 256-288 | 0.0588 |
| 23 | SSC | V27 | 832-864 | 0.0493 | FSC | V52 | 608-640 | 0.0596 | CD15 | V69 | 128-160 | 0.0760 | CD34 | V106 | 288-320 | 0.0474 |
| 24 | SSC | V17 | 512-544 | 0.0483 | FSC | V63 | 960-992 | 0.0571 | CD15 | V65 | 0-32 | 0.0712 | CD34 | V124 | 864-896 | 0.0426 |
| 25 | SSC | V28 | 864-896 | 0.0462 | FSC | V53 | 640-672 | 0.0520 | CD15 | V87 | 704-736 | 0.0696 | CD34 | V104 | 224-256 | 0.0422 |
| 26 | SSC | V5 | 128-160 | 0.0270 | FSC | V54 | 672-704 | 0.0507 | CD15 | V91 | 832-864 | 0.0541 | CD34 | V122 | 800-832 | 0.0272 |
| 27 | SSC | V6 | 160-192 | 0.0252 | FSC | V45 | 384-416 | 0.0288 | CD15 | V90 | 800-832 | 0.0516 | CD34 | V128 | 992-1024 | 0.0265 |
| 28 | SSC | V3 | 64-96 | 0.0174 | FSC | V43 | 320-352 | 0.0195 | CD15 | V95 | 960-992 | 0.0276 | CD34 | V121 | 768-800 | 0.0252 |
| 29 | SSC | V2 | 32-64 | 0.0125 | FSC | V33 | 0-32 | 0.0000 | CD15 | V93 | 896-928 | 0.0241 | CD34 | V127 | 960-992 | 0.0218 |
| 30 | SSC | V31 | 960-992 | 0.0093 | FSC | V34 | 32-64 | 0.0000 | CD15 | V96 | 992-1024 | 0.0198 | CD34 | V125 | 896-928 | 0.0212 |
| 31 | SSC | V1 | 0-32 | 0.0000 | FSC | V35 | 64-96 | 0.0000 | CD15 | V92 | 864-896 | 0.0154 | CD34 | V123 | 832-864 | 0.0211 |
| 32 | SSC | V4 | 96-128 | 0.0000 | FSC | V36 | 96-128 | 0.0000 | CD15 | V94 | 928-960 | 0.0129 | CD34 | V126 | 928-960 | 0.0181 |

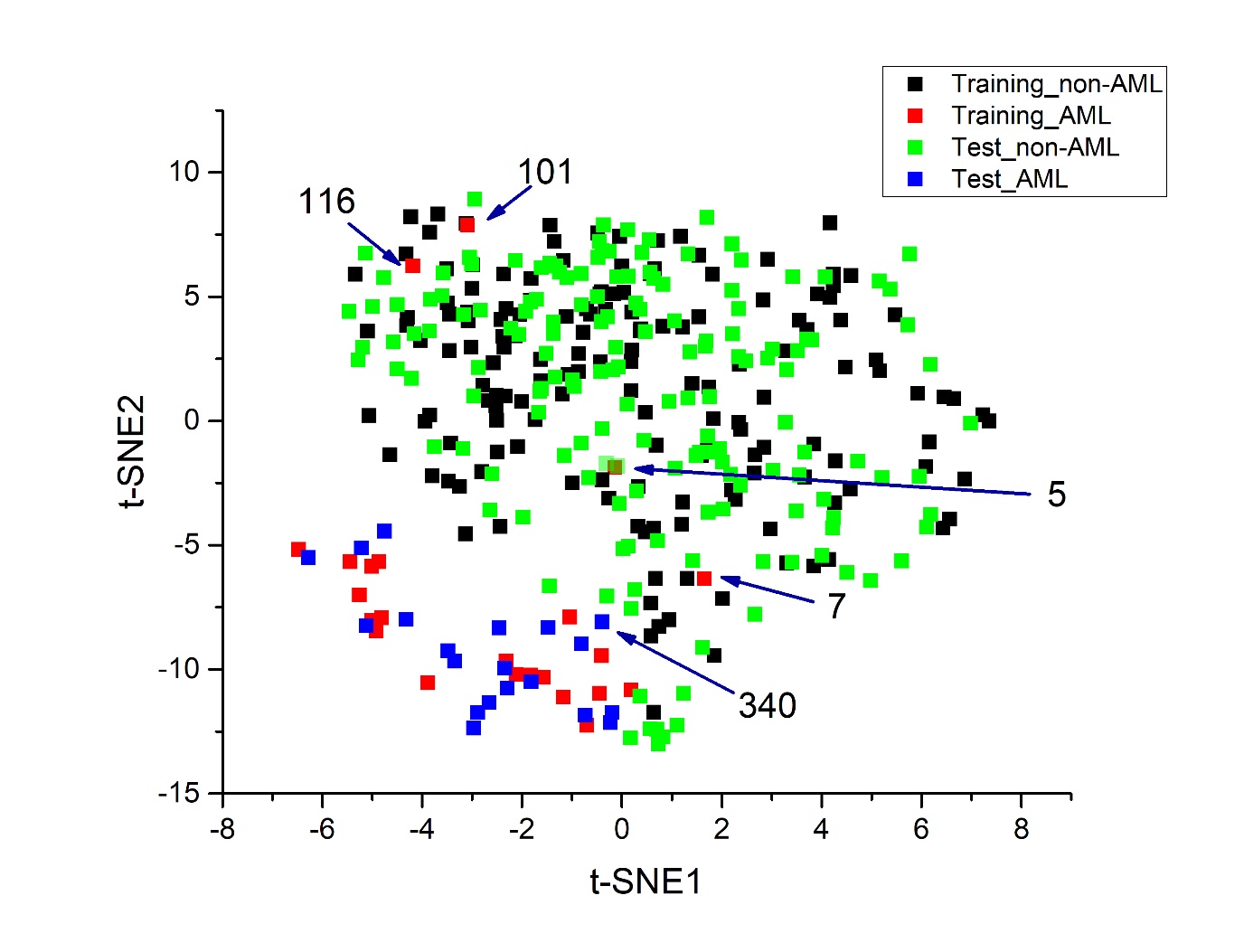

S1 Fig. 2D t-SNE plot for training and test set

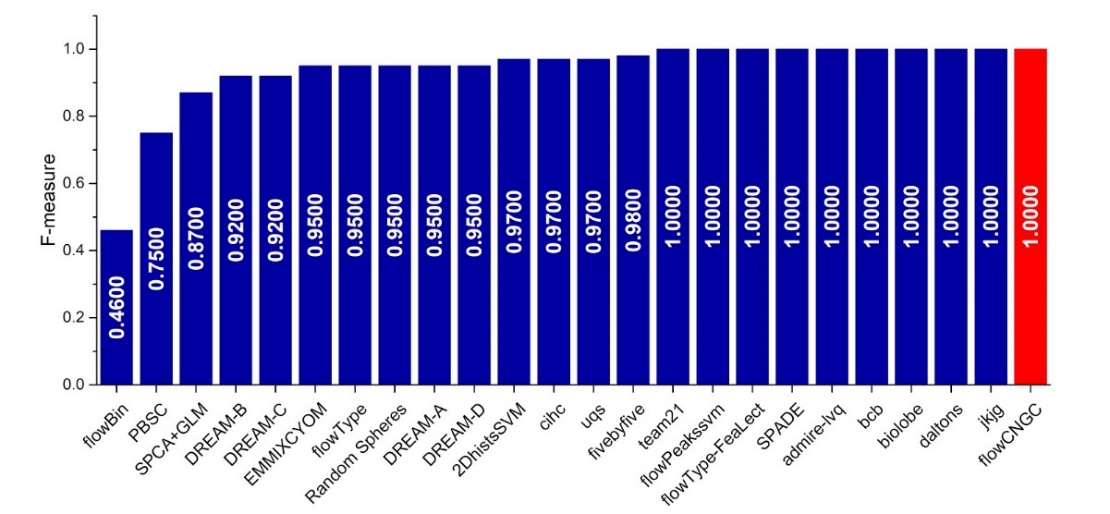

S2 Fig. Model performance comparison with algorithms used in FlowCAP-II Challenge
